# Rapid Genomic Characterization and Global Surveillance of *Klebsiella* Using Pathogenwatch

**DOI:** 10.1101/2021.06.22.448967

**Authors:** Silvia Argimón, Sophia David, Anthony Underwood, Monica Abrudan, Nicole E. Wheeler, Mihir Kekre, Khalil Abudahab, Corin A. Yeats, Richard Goater, Ben Taylor, Harry Harste, Dawn Muddyman, Edward J. Feil, Sylvain Brisse, Kathryn Holt, Pilar Donado-Godoy, KL Ravikumar, Iruka N. Okeke, Celia Carlos, David M. Aanensen, The NIHR Global Health Research Unit for the Genomic Surveillance of Antimicrobial Resistance

## Abstract

**Background:** *Klebsiella* species, including the notable pathogen *K. pneumoniae*, are increasingly associated with antimicrobial resistance (AMR). Genome-based surveillance can inform interventions aimed at controlling AMR. However, its widespread implementation requires tools to streamline bioinformatic analyses and public health reporting.

**Methods:** We developed the web application Pathogenwatch, which implements analytics tailored to *Klebsiella* species for integration and visualization of genomic and epidemiological data. We populated Pathogenwatch with 16,537 public *Klebsiella* genomes to enable contextualization of user genomes. We demonstrated its features with 1,636 genomes from four low- and middle-income countries (LMICs) participating in the NIHR Global Health Research Unit (GHRU) on AMR.

**Results:** Using Pathogenwatch, we found that GHRU genomes were dominated by a small number of epidemic drug-resistant clones of *K. pneumoniae*. However, differences in their distribution were observed (e.g. ST258/512 dominated in Colombia, ST231 in India, ST307 in Nigeria, ST147 in the Philippines). Phylogenetic analyses including public genomes for contextualization enabled retrospective monitoring of their spread. In particular, we identified hospital outbreaks, detected introductions from abroad, and uncovered clonal expansions associated with resistance and virulence genes. Assessment of loci encoding O-antigens and capsule in *K. pneumoniae*, which represent possible vaccine candidates, showed that three O-types (O1-O3) represented 88.9% of all genomes, whereas capsule types were much more diverse.

**Conclusions:** Pathogenwatch provides a free, accessible platform for real-time analysis of *Klebsiella* genomes to aid surveillance at local, national and global levels. We have improved representation of genomes from GHRU participant countries, further facilitating ongoing surveillance.

**summary:** Pathogenwatch is a free web-application for analysis of *Klebsiella* genomes to aid surveillance at local, national and global levels. We improved the representation of genomes from middle-income countries through the Global Health Research Unit on AMR, further facilitating ongoing surveillance.

## INTRODUCTION

The *Klebsiella* genus, which belongs to the *Enterobacteriaceae* family, comprises several species that cause opportunistic infections in hospital and community settings [1, 2]. *Klebsiella* species are found in the environment, and commonly contaminate healthcare environments and medical equipment [3, 4]. They also frequently colonize the intestinal tract and other mucosal surfaces of humans, which can serve as reservoirs for infection [1, 5, 6]. Infections occur most commonly in the elderly, neonates and in immunocompromised individuals, and include urinary tract infections, pneumonia, bloodstream infections and sepsis [7, 8]. By far the most clinically significant member of the genus is *Klebsiella pneumoniae* [9]. However, other species including *K. quasipneumoniae*, *K. variicola*, and *K. oxytoca* are also notable pathogens [10–12].

In recent years, the prevalence of infections caused by *K. pneumoniae* that are multi-drug resistant has risen sharply [13]. Increasing resistance levels have largely been driven by the emergence of strains producing extended spectrum beta-lactamase (ESBL) and carbapenemase enzymes, which are typically plasmid-encoded. ESBLs confer resistance to third-generation cephalosporins and monobactams, while carbapenemases result in resistance to almost all beta-lactams including carbapenems [14, 15]. In the community, hypervirulent *K. pneumoniae* can cause severe infections owing to the production of specific virulence factors including siderophores and the capsule expression regulator RmpA [16]. Moreover, infections and outbreaks involving strains with both multi-drug resistance and hypervirulence have now also been reported, leading to concern over a potential rise of serious untreatable infections [17, 18].

Increasing constraints around treatment options for multidrug-resistant *Klebsiella* infections, and accompanying rise in hypervirulence, have led to an urgent need for novel drugs. Some have reached the market recently, including ceftazidime-avibactam and plazomicin, but more are needed. There is also renewed interest in the development of a preventative vaccine for *K. pneumoniae* infections [19, 20]. Potential vaccine candidates include K-antigens belonging to the bacterial capsule polysaccharide (CPS) and O-antigens comprising the outermost part of the lipopolysaccharide (LPS) [21]. However, these antigens are variable and may differ across geographic regions or infection types, underlining the need for seroepidemiology.

Pathogen surveillance provides a powerful tool for understanding the evolution and spread of resistant bacteria and defining their clinically relevant features [22]. With more widespread adoption of genomic approaches, there is a growing need for tools that streamline whole-genome sequencing (WGS) analyses, circumvent the need for user expertise in bioinformatics, and deliver results for public health utility. Here we describe the *Klebsiella* scheme of the web application Pathogenwatch, which incorporates community-driven tools together with additional functionality to provide detailed typing and phylogenetic analyses of *Klebsiella* isolates, and integration of genomic and epidemiological data [23, 24]. We illustrate features of Pathogenwatch by analyzing *Klebsiella* genomes from four LMICs participating in the National Institute for Health Research Global Health Research Unit (GHRU) on genomic surveillance of AMR. We also demonstrate how it can aid decision making in real time at local and global scales and inform the choice and effectiveness of key interventions such as vaccines.

## METHODS

### Assembly and curation of public *Klebsiella* genomes for Pathogenwatch

We identified 18 319 samples in the European Nucleotide Archive (ENA) labelled either “*Klebsiella*” or “*Raoultella*” (a closely related genus that is not phylogenetically separate from *Klebsiella*, henceforth included within “*Klebsiella*”) with paired-end Illumina sequence data and geolocation data, as of August 3, 2020. *De novo* assembly using the raw sequence data was attempted using a SPAdes pipeline, resulting in the assembly of 97.1% (17 783/18 319) of the samples [25]. Various quality control (QC) metrics were used to discard assemblies of poor quality (Supplementary Tables 1 and 2). 93.0% (16 537/17 783) of assemblies passed these QC criteria and were imported into Pathogenwatch as public genomes. Metadata for these samples (Supplementary Table 3) were downloaded via the ENA application programming interface (API), curated, and linked to the assemblies in Pathogenwatch.

### Pathogenwatch features tailored to *Klebsiella* species

#### Species determination

The Speciator tool assigns species by comparing assemblies to genomes within a reference library via Mash [26, 27]. This library comprises a manually curated set of reference assemblies tailored for *Klebsiella* and other *Enterobacteriaceae* species from Kleborate (currently v. 2.0.1) [28].

#### Genomic characterization

Multi-locus sequence typing (MLST) and core genome MLST (cgMLST) are performed using the allelic and profile definitions from databases hosted via the BIGSdb platform at Institut Pasteur [29]. Resistance and virulence loci, K- and O-loci, and *wzi* genes are typed in Pathogenwatch via an implementation of Kleborate (currently v. 2.0.1). The *wzi* genes are defined using the BIGSdb-Pasteur platform [30]. Plasmid replicons are identified using Inctyper (currently v.0.0.4), with the *Enterobacteriaceae* database (5/11/2020 version) from PlasmidFinder [31, 32].

#### Phylogenetic analyses

The pan-genome tool Roary was used previously to identify 2539 genes present in ≥95% of genomes from each species within a European collection of *K. pneumoniae* species complex isolates (*K. pneumoniae*, *K. quasipneumoniae*, *K. variicola*, *K. quasivariicola*) [22, 33]. To further define a set of core genes in *K. pneumoniae* for phylogenetic analyses within Pathogenwatch, we first merged any matches to the 2539 genes that were overlapping in any of 13 diverse *K. pneumoniae* reference genomes to form pseudo-sequences (Supplementary Table 4). We then removed any genes/pseudo-sequences that were paralogous (i.e. matched another with >80% nucleotide identity and an e-value of <1e-35 via BLASTn) and any that were absent or incomplete in one or more references [34]. The core gene library used in Pathogenwatch was thus comprised of the remaining 1972 genes (or pseudo-sequences). These are queried to generate pairwise SNP distances between genomes, which are used to construct neighbor-joining trees.

### Whole genome sequencing and assembly of GHRU isolates

Laboratories in four participant countries of the GHRU obtained 1706 *Klebsiella* genomes from isolates collected in 2013-2019 (Colombia, *n*=589; India, *n*=347; Nigeria, *n*=164; Philippines, *n*=606) [35–38]. Of the samples from the Philippines, 342/606 (56.4%) have been previously reported [39]. Raw sequence data were assembled and quality-checked as described in this Methods section. 96.0% (1636/1706) of assemblies passed QC and were analyzed with Pathogenwatch (Supplementary Tables 5 and 6). Of these 1636 isolates, 1625 were from human clinical samples and 11 from environmental samples (Nigeria only). Raw data have been deposited in the ENA under study accessions ERP112087, ERP112088, ERP112089, ERP112091, and ERP019480.

## RESULTS & DISCUSSION

### User upload and characterization of *Klebsiella* genomes using Pathogenwatch

Here we present features of the Pathogenwatch application (https://pathogen.watch/) that are tailored to genomic analysis of *Klebsiella* (including the closely-related *Raoultella* genus). Users can upload either assemblies or raw sequence reads to Pathogenwatch, the latter of which will be assembled via a SPAdes pipeline [25]. Assemblies identified by Pathogenwatch as belonging to *Klebsiella* are then subjected to specific analytic pipelines (Figure 1). These include MLST and cgMLST for species with available schemes, identification of resistance genes, virulence loci, capsule and O-antigen biosynthesis loci, and replicon typing (Supplementary Table 7).

**Figure 1.**
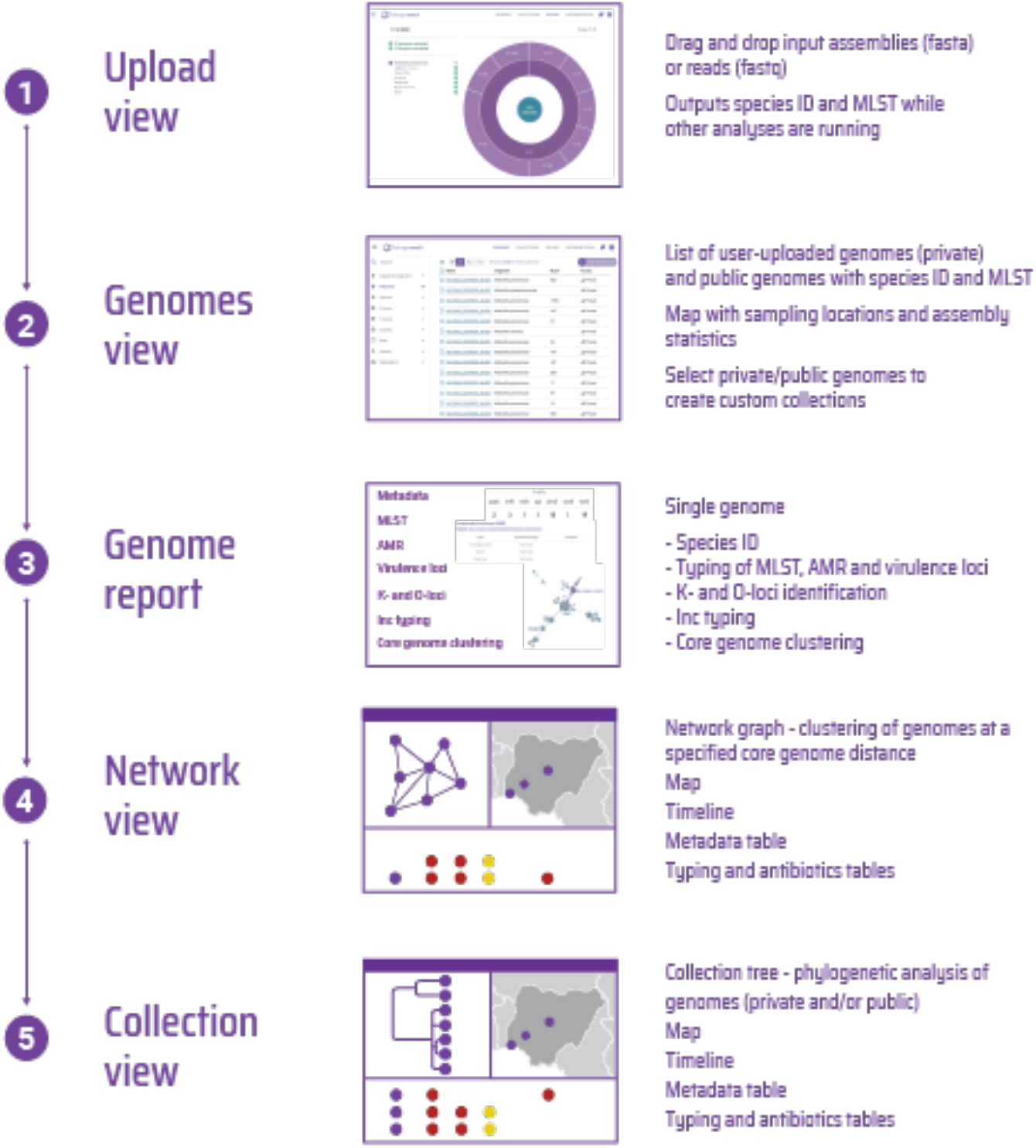
Overview of the analytical processes performed on *Klebsiella* genomes and the available visualizations in Pathogenwatch.

**Figure 2.**
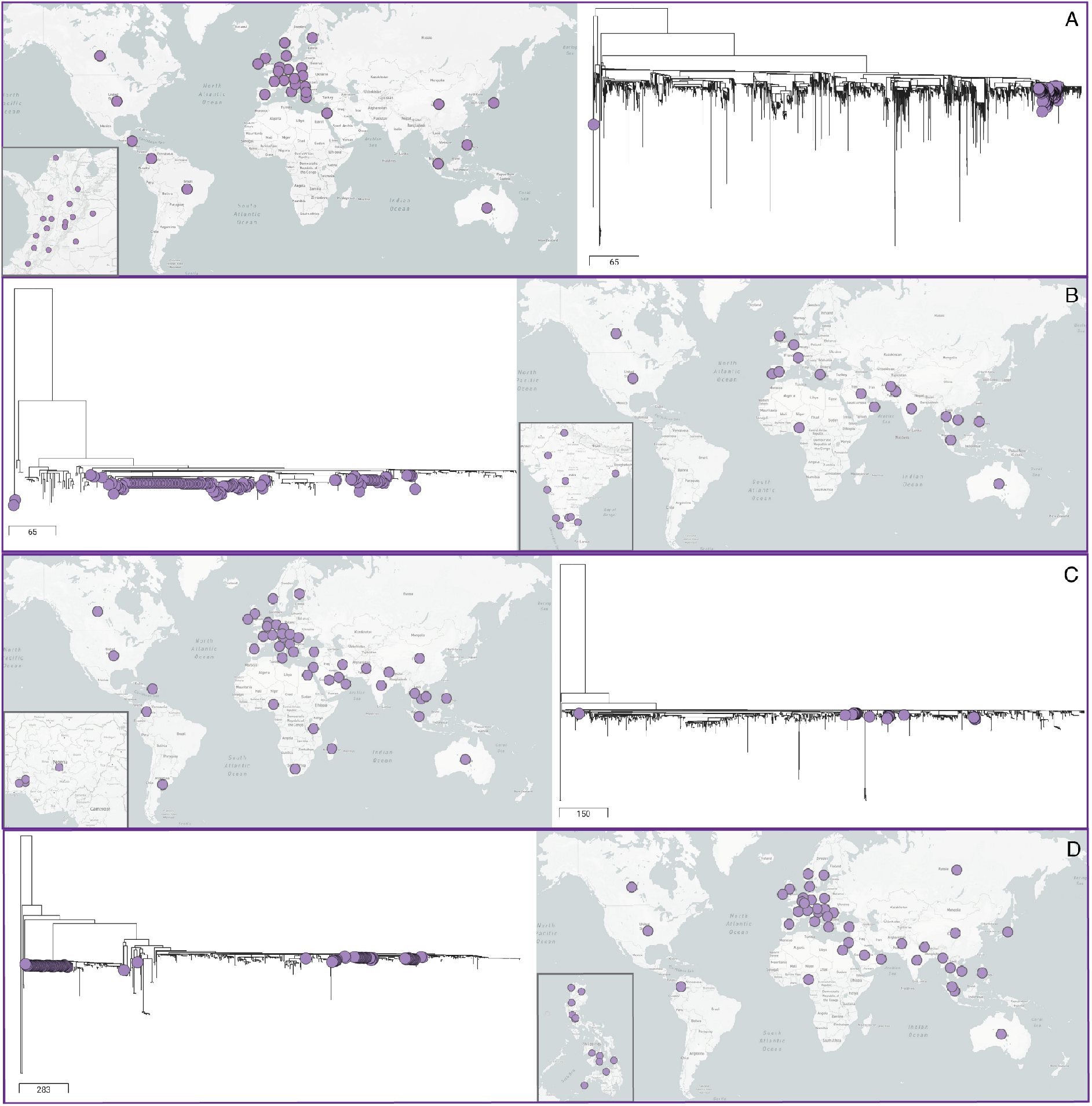
Pathogenwatch shows different dynamics of transmission and dissemination of the dominant “high-risk” lineages in each GHRU participant country. ST258 genomes from Colombia form one main phylogenetic cluster, suggestive of a single successful introduction (**A**). ST231 genomes from India (**B**), ST307 genomes from Nigeria (**C**), and ST147 genomes from the Philippines (**D**), all form multiple phylogenetic clusters, suggesting multiple origins. The map insets show the widespread distribution of these clones in each country.

As standard in Pathogenwatch, users can browse public and/or uploaded genomes. Public genomes include 16 537 high-quality *Klebsiella* genomes with geolocation data (Supplementary Table 3; Supplementary Figure 1). All metadata and results from the analytic pipelines can be viewed and downloaded for an individual genome in a “Genome report” or collectively for multiple selected genomes from the “Genomes” page. Within the Genome report, a clustering tool can be used to rapidly identify the most closely-related genomes (from all public and uploaded genomes) to a genome of interest based on cgMLST allelic differences. Users can generate an interactive network visualization of genomes clustered within a particular allelic threshold.

We have also developed the ability for users to generate a phylogenetic tree comprising multiple selected genomes of *K. pneumoniae* (comprising user and/or public genomes). The tree is functionally integrated in the “Collection” view with a map and timeline, showing the locations and sampling dates of genomes if provided, and results from all analytic pipelines. This visualization enables the user to interactively explore the data, while the tree and all other data from individual collections can be downloaded in standard formats.

All sequence data and metadata uploaded by users remain private to their accounts. Genomes grouped into collections are also kept private by default, although they can be shared with collaborators via a URL. There is also an option for users to integrate confidential metadata into visualizations locally within the browser, without upload of data to the Pathogenwatch server.

Pathogenwatch, as well as the integrated tools, is being actively developed and will be updated periodically to provide the latest typing information (including new resistance and virulence mechanisms), newly-available public genomes, and other features. The modular architecture of Pathogenwatch also enables integration of new analytics. Detailed descriptions of all of the above processes can be found in the documentation [40].

Below we describe the utility of Pathogenwatch for epidemiological surveillance with 1636 isolates collected from four laboratories linked to national or country-wide networks in Colombia, India, Nigeria, and the Philippines (Supplementary Table 6) [41]. All four countries have been previously underrepresented in genomic surveillance efforts (Table 1) despite previous estimates of significant burden of *Klebsiella* infections in those regions [42].

**Table 1.**
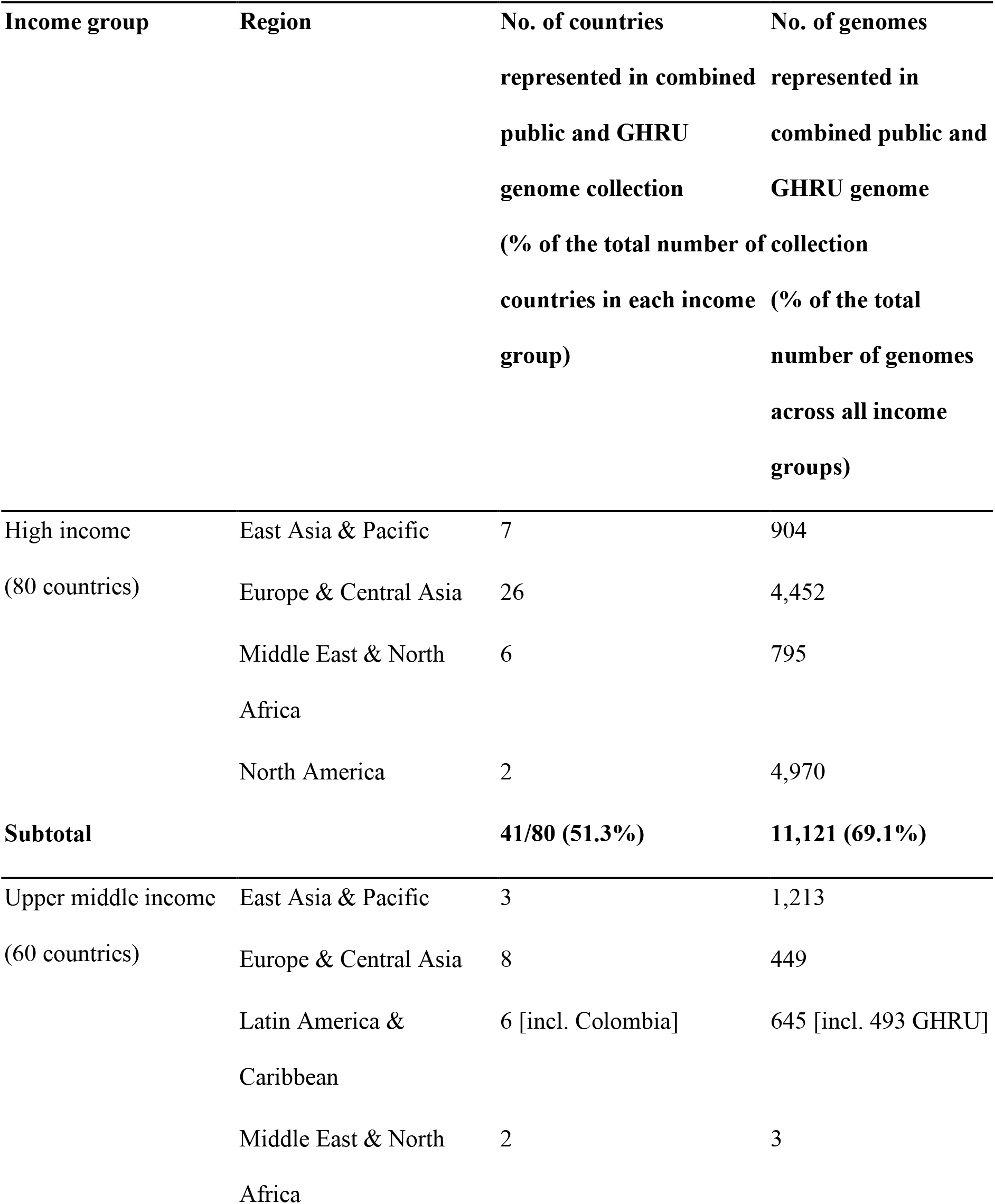
Distribution of *K. pneumoniae* genomes in the combined public and GHRU collection by region and country income class.

### Species identification

Pathogenwatch first assigns genome assemblies to a species via the Speciator tool. The assigned species then determines the downstream analyses. Speciator can currently identify assemblies belonging to *K. pneumoniae*, *K. quasipneumoniae*, *K. variicola*, *K. quasivariicola* and *K. Africana* (which together make up the *K. pneumoniae* species complex), as well as 11 other *Klebsiella* species (Supplementary Table 7).

*K.pneumoniae* accounted for 88.5% (14 635/16 537) and 88.7% (1 451/1 636) of *Klebsiella* genomes from the public and GHRU collections, respectively (Supplementary Table 8), reaffirming the clinical dominance of this species. The other most frequently observed species were *K. quasipneumoniae*, *K. variicola*, *K. aerogenes*, and *K. michiganensis* (comprising 4.0%, 1.9%, 1.8%, and 1.8% of the combined collections, respectively). As shown previously by others, we found inaccuracies in laboratory identification methods for *Klebsiella* [43]. For example, of the 1576 isolates in the GHRU collection assigned to *K. pneumoniae* using laboratory methods, Speciator identified 147 (9.3%) as *K. quasipneumoniae*, 4 (0.3%) as *K. variicola*, 2 (0.1%) as *K. michiganensis*, 2 (0.1%) as *K. oxytoca*, and 2 (0.1%) as *K. quasivariicola* (Supplementary Table 9).

### Surveillance of high-risk clones

Assemblies identified by Pathogenwatch as *Klebsiella* are subject to MLST and/or cgMLST based on the availability of schemes (Supplementary Table 7). The majority of our GHRU *K. pneumoniae* genomes belonged to a small number of known epidemic (“high-risk”) sequence types (ST) that were also over-represented in the public genome collection. In particular, 51.7% (750/1451) of the GHRU genomes and 56.7% (8295/14 635) public genomes belonged to only 10 STs that were the most frequently observed across the combined collections (Table 2). Overall, a high number of STs were observed in both collections (209 and 1115 in the GHRU and public collections, respectively). We also found 62 STs present among GHRU genomes that were not identified among public genomes, of which 33 were novel.

**Table 2.**
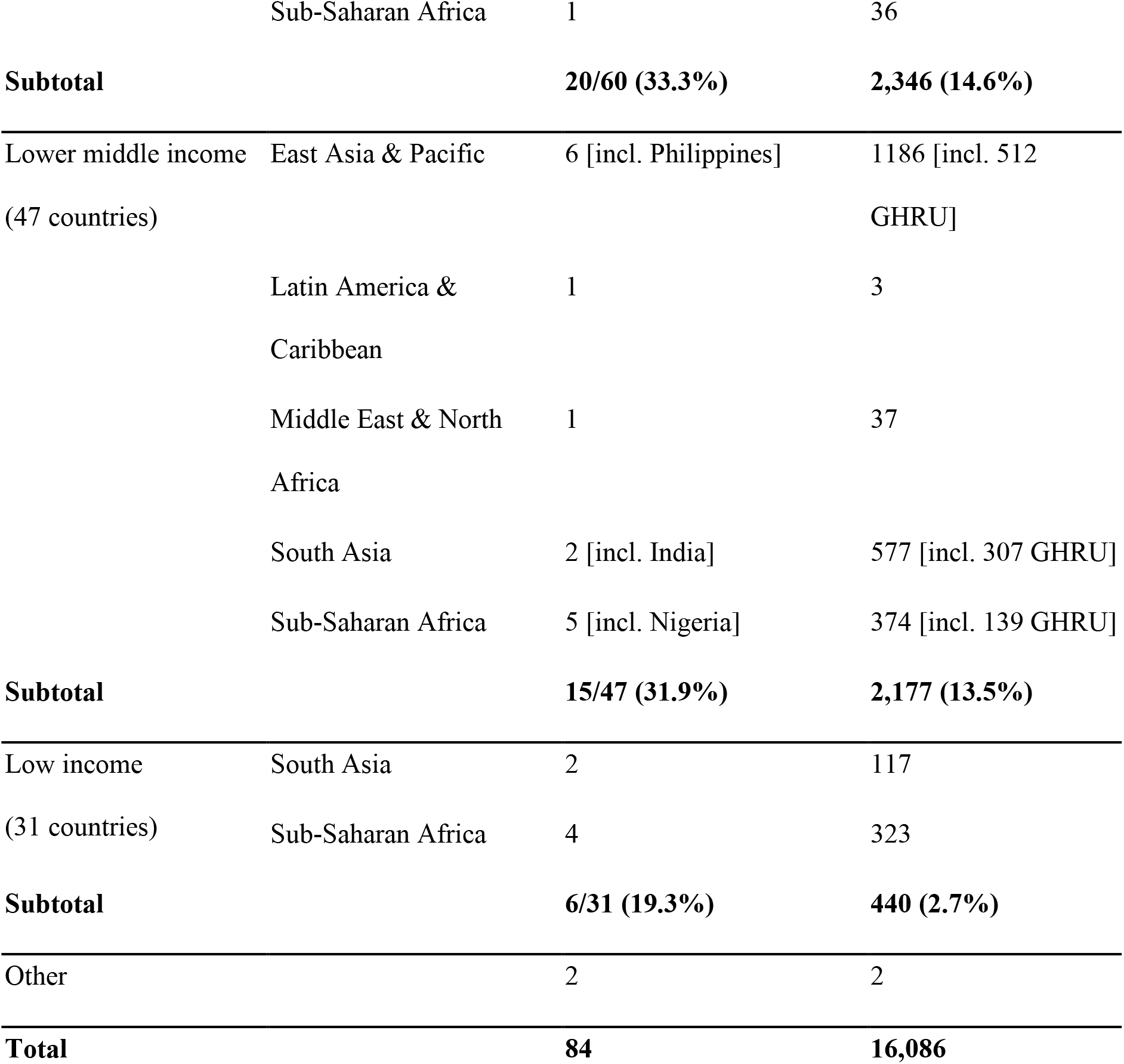
Characteristics of the top ten most frequently-observed sequence types (ST) of *K. pneumoniae* in the combined public and GHRU genome collections. Provided in Excel document.

Clonal lineages of *K. pneumoniae* differ in their ability to acquire resistance and virulence genes, and in their propensity to spread within hospital and community environments [44]. Closing geographic gaps in genomic surveillance to fully describe the diversity of clones circulating across different regions is therefore vital for a better understanding of the local epidemiology of *K. pneumoniae* infections. We found clear differences in the dominant high-risk STs of *K. pneumoniae* circulating in the GHRU countries, with the single largest contribution from ST258 in Colombia (24.3%), ST231 in India (34.5%), ST307 in Nigeria (15.1%), and ST147 in the Philippines (20.1%) (Supplementary Figure 2). Each of these STs is widely disseminated across each country. Despite differences in sampling strategies between countries, these results are in line with previous observations relating to the regional distribution of high-risk STs [43–48].

The ability of Pathogenwatch to generate phylogenetic trees of *K. pneumoniae* linked to metadata and other analytics provides a platform for monitoring the spread of lineages, offering relevant insights at global through to local scales. For example, ST258 genomes from Colombia form a single major cluster within the global phylogeny, which represents isolates from 31 countries (Figure 1A). This indicates that a single main introduction followed by within-country spread is likely responsible for the high endemicity of this lineage reported in Colombia [49]. By contrast, multiple phylogenetic clusters of ST231, ST307, and ST147 are observed in India, Nigeria, and the Philippines, respectively, demonstrating different origins for the multiple circulating lineages (Figure 1B–D). At a local scale, we found evidence of clonal spread of OXA-181- and CTX-M-15-producing ST147 within a hospital in India, over a period of three years (Supplementary Figure 3). However, some ST147 isolates collected from the same hospital during this period were phylogenetically distinct from the outbreak cluster, and thus the respective patients could be ruled out of the outbreak.

### Detection of resistance and virulence mechanisms

Known resistance and virulence loci are identified in Pathogenwatch via Kleborate [28]. Resistance mechanisms currently include SNPs, acquired genes, and gene truncations that are relevant for different antibiotics or antibiotic classes. Virulence genes include those encoding acquired siderophores (yersiniabactin, salmochelin, aerobactin), the genotoxin colibactin, the hypermucoidy locus *rmpADC*, and alternative hypermucoidy marker gene *rmpA2*.

A high proportion of GHRU *K. pneumoniae* isolates contained an ESBL or carbapenemase gene (75.2% and 63.0%, respectively). This was also seen in the public genomes, with 54.5% (7983/14 635) of *K. pneumoniae* genomes carrying an ESBL, and 57.5% (8416/14 635) carrying a carbapenemase. These rates far exceed those observed in most clinical settings to date (i.e. typically <20% for carbapenemase-producing isolates) and demonstrate the tendency to prioritize multi-drug resistant isolates for sequencing [39, 50–52].

Despite sampling biases, clear differences existed between *K. pneumoniae* genomes from the GHRU countries with regards to major carbapenemase genes circulating (e.g. KPC genes dominate in Colombia, NDM genes in the Philippines and Nigeria, OXA-48-like genes in India (Supplementary Figure 4)). These findings are in line with broader regional patterns reported previously and also uncovered using the public genomes [53–56]. In contrast, CTX-M-15 was consistently the most frequently observed ESBL gene, carried by 72.9-100% of ESBL-producing *K. pneumoniae* in each country.

Assessment of the prevalence and dissemination of mobile colistin resistance (*mcr*) genes, the first variant of which was discovered in 2015, among *K. pneumoniae* genomes in the public and GHRU collections revealed that these are still rare across all regions [57]. Only 0.3% (5/1636) of GHRU genomes carried an *mcr* gene, and 0.9% (128/14 635) of public genomes did. Among the latter, 52.3% (67/128) carried *mcr-9*.

We found that the majority (>80%) of all *K. pneumoniae* genomes in both public and GHRU genome collections had either no known acquired virulence factors or yersiniabactin only (Supplementary Table 10). However, there was an over-representation of colibactin among *K. pneumoniae* GHRU genomes from Colombia (present in 24.7% of isolates), which was associated with ST258. Furthermore, 37.5% of *K. pneumoniae* GHRU genomes from India carried aerobactin, which was almost always carried by ST231 or ST2096. In particular, phylogenetic analysis using a collection of ST231 isolates from both the GHRU and public collections highlighted a sublineage that has acquired aerobactin and yersiniabactin, as well as the OXA-232 carbapenemase (Figure 3A; see also [36]).This convergence of both resistance and virulence has been coupled with rapid clonal expansion and international spread, and close monitoring is needed.

**Figure 3.**
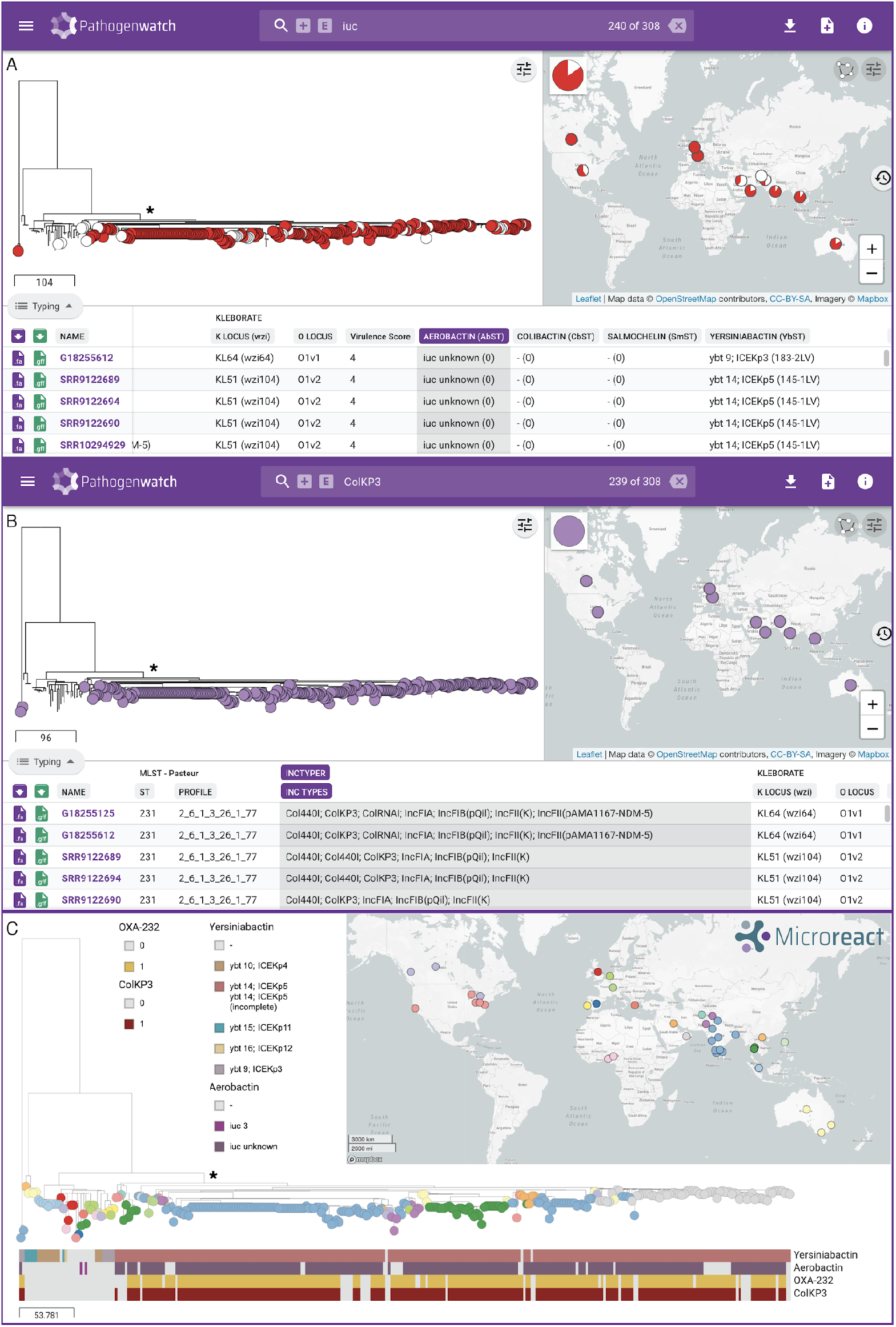
Pathogenwatch demonstrates convergence of virulence and resistance in a phylogenetic tree of 308 ST231 genomes from the public and GHRU collections (clade indicated with *). **(A)** The tree and map are filtered via the search bar by the presence of the virulence determinant aerobactin (*iuc*). All aerobactin-positive isolates are indicated with a circular node in the tree (red or white). Red nodes indicate the additional presence of the OXA-232 carbapenemase gene. Pie charts on the map show the relative proportion of aerobactin-positive isolates with and without OXA-232 **(B)** The tree and map are filtered via the search bar by the presence of replicon sequence ColKP3. ColKP3-positive isolates are indicated with purple nodes in the tree. **(C)** Likely acquisition of virulence loci (yersiniabactin and aerobactin) and plasmid-borne resistance (OXA-232 and ColKP3) followed by clonal expansion of the clade indicated with (*).

### Monitoring of mobile genetic elements

Plasmid replicons in any *Klebsiella* genome are identified in Pathogenwatch using the Inctyper tool. While it is not usually possible to directly link resistance or virulence genes to particular plasmids with short-read assemblies, we can nevertheless gain important epidemiological insights by analyzing patterns in the diversity and distribution of plasmid replicons.

For example, we noted that 68.7% (211/307) of GHRU *K. pneumoniae* isolates from India carry the ColKP3 replicon, which was not found in isolates from any of the other three countries. This replicon was previously found in a conserved 6.1kb ColE-type plasmid initially reported from or linked to international travel to India but since identified elsewhere in patients without travel history and causing local outbreaks [59–62]. Of the 211 isolates with a ColKP3 replicon, all but one carry either the OXA-232 (*n*=168, 79.6%) or OXA-181 (*n*=42, 19.9%) carbapenemase.

Using the combined collection of GHRU and public *K. pneumoniae* genomes, we confirmed a strong association between the ColKP3 plasmid and the OXA-232 gene. In particular, we found that 86.7% (684/789) of *K. pneumoniae* isolates with a ColKP3 plasmid possess an OXA-232 gene, compared to 0.2% (35/15 297) of those without (Pearson’s Chi-square = 13 136.39, p<0.0001). The association between ColKP3 and OXA-232 does not appear to be an artefact of lineage or geographic effects, as we found ColKP3/OXA-232 isolates in 45 different STs and 18 different countries overall. However, the majority (79.7%) do belong to only four STs (14, 16, 231, 2096), and they originate mostly from South and Southeast Asia, and the Arabian Peninsula. It has previously been suggested that disruption of the IS*Ecp1* transposase may have stabilized the OXA-232 gene on the ColKP3 plasmid [59].

Phylogenetic analysis of all ST231 isolates in Pathogenwatch suggested that the ColKP3 plasmid was acquired once, and then disseminated vertically through the lineage via clonal spread (Figure 3–C) (see also [36]).

### K- and O-loci monitoring to aid vaccine development

O-antigen biosynthesis loci (O-loci) and capsular loci (including the *wzi* alleles and K-loci) present in *Klebsiella* genomes are identified in Pathogenwatch via Kleborate. Studies have reported development of a *K. pneumoniae* (and *P. aeruginosa*) glycoconjugate vaccine based on the O-serotypes O1, O2, O3, and O5, and another for hypervirulent *K. pneumoniae* based on the K1 and K2 capsule types [19, 20]. It is thus crucial to monitor the diversity of O- and K-types across different lineages, geographic regions, age groups, clinical sources, and over time, to ensure that a potential vaccine will adequately protect target populations.

Despite the biases present in both the public and GHRU sample collections, the breadth of geographic representation and inclusion of countries previously under-represented make it a valuable collection to describe the diversity of O- and K-types. Here we considered human-associated isolates from the combined public and GHRU genomes which had O-types and K-types assigned with a confidence level of “good” or better by Kleborate.

We found that the O1, O2, and O3 serotypes (including their subtypes) were the most prevalent, comprising 88.9% (10 252/11 530) of *K. pneumoniae* isolates, and in line with previous reports [21]. Major high-risk STs with high levels of multidrug resistance were also dominated by these serotypes (Table 2). Other serotypes present in >1% of *K. pneumoniae* isolates included O4 (5.6%), OL101 (2.7%), and O5 (1.9%). As vaccines may be developed to target high-risk populations such as neonates, we also stratified the O-types identified in the GHRU isolates by patient age. We found that O1, O2, and O3 represented 52.9-91.4% of *K. pneumoniae* isolates from each age group (Supplementary Figure 5). Furthermore, we noted that the distribution of O-types varied substantially across species. For example, despite dominating in *K. pneumoniae* (88.9%), O1-O3 together made up only 40.5% (145/358) and 49.7% (94/189) of isolates from *K. quasipneumoniae* and *K. variicola*, respectively. Meanwhile, O5 was far more prevalent in both species (found in 29.9% and 40.7% isolates, respectively) than in *K. pneumoniae* (1.9%).

In contrast with the O-types, the 5 most common K-loci (KL) types (KL107, KL106, KL102, KL64, and KL51) represented only 47.8% (5218/10 922) of *K. pneumoniae* isolates, and a minimum of 39 KL-types were required to encompass ≥90% of genomes. High-risk multidrug-resistant lineages were typically dominated by just one or two KL-types, although high numbers of types in some lineages also illustrate the capacity for genetic exchange of the *cps* locus. We found that the most frequently observed K-loci were not consistently present across all age groups sampled in the GHRU collection (Supplementary Figure 5), although larger sample collections of target populations with consistent sampling will be required for a robust assessment.

Likewise, future analyses using representative sample collections from target populations will further elucidate key trends and differences in K- and O-loci diversity. The importance of combining patient information with genomic data for developing insights relevant to patient outcomes cannot be understated. The ability of Pathogenwatch to combine these data with the temporal and spatial trends of clonal lineages, and multidrug resistance and virulence, provides a rational system for informing the development of vaccines and therapeutics, and also for monitoring population changes as a consequence of implementing interventions.

## CONCLUDING REMARKS

WGS empowers AMR surveillance laboratories to make public health decisions by providing a high-resolution view of the circulating bacterial strains and aiding outbreak investigations. Here we have presented the features of Pathogenwatch, a free, accessible platform for characterization and contextualization of *Klebsiella* genomes to aid surveillance at local, national and global levels. The newly-built capacity and expertise of four laboratories in LMICs to undertake ongoing genome sequencing, as developed during the wider GHRU project, will be enhanced by the use of Pathogenwatch and the increased representation of genomes from their countries. Extending this model to laboratories in other LMICs is a future priority.

## ACKNOWLEDGMENTS

Members of the NIHR Global Health Research Unit on Genomic Surveillance of Antimicrobial Resistance: Johan Fabian Bernal, Alejandra Arevalo, Maria Fernanda Valencia, and Erik C. D. Osma Castro of the Colombian Integrated Program for Antimicrobial Resistance Surveillance – Coipars, CI Tibaitatá, Corporación Colombiana de Investigación Agropecuaria (AGROSAVIA), Tibaitatá – Mosquera, Cundinamarca, Colombia; Geetha Nagaraj, Varun Shamanna, Vandana Govindan, Akshata Prabhu, D. Sravani, M. R. Shincy, Steffimole Rose, and Ravishankar K.N of the Central Research Laboratory, Kempegowda Institute of Medical Sciences, Bengaluru, India; Anderson O. Oaikhena, Ayorinde O. Afolayan, Jolaade J Ajiboye, and Erkison Ewomazino Odih of the Department of Pharmaceutical Microbiology, Faculty of Pharmacy, University of Ibadan, Oyo State, Nigeria; Marietta L. Lagrada, Polle Krystle V. Macaranas, Agnettah M. Olorosa, June M. Gayeta, Melissa Ana L. Masim, and Elmer M. Herrera of the Antimicrobial Resistance Surveillance Reference Laboratory, Research Institute for Tropical Medicine, Muntinlupa, the Philippines; Ali Molloy, alimolloy.com; John Stelling, The Brigham and Women’s Hospital; and Carolin Vegvari, Imperial College London.

We are grateful to the DNA Pipelines and Pathogen Informatics teams at the Wellcome Sanger Institute for their support.

## FUNDING

This work was supported by Official Development Assistance (ODA) funding from the National Institute of Health Research [grant number 16_136_111].

This research was commissioned by the National Institute of Health Research using Official Development Assistance (ODA) funding. The views expressed in this publication are those of the authors and not necessarily those of the NHS, the National Institute for Health Research or the Department of Health.

## CONFLICT OF INTEREST

The authors: No reported conflicts of interest. All authors have submitted the ICMJE Form for Disclosure of Potential Conflicts of Interest.

## ABBREVIATIONS

AMR: antimicrobial resistance
LMICs: low- and middle-income countries
GHRU: Global Health Research Unit
ST: sequence type
ESBL: extended-spectrum beta-lactamase
CPS: capsular polysaccharide
LPS: lipopolysaccharide
WGS: whole genome sequencing
ENA: European Nucleotide Archive
QC: quality control
API: application programming interface
MLST: multi-locus sequence typing
cgMLST: core genome multi-locus sequence typing
SNP: single nucleotide polymorphism
KL: K-loci

## Notes

### Competing Interest Statement

The authors have declared no competing interest.

