## Supplementary Information for "Rapid Genomic Characterization and Global Surveillance of *Klebsiella* Using Pathogenwatch"

[a] Members of the NIHR Global Health Research Unit on Genomic Surveillance of Antimicrobial Resistance are listed in the Acknowledgments.

#### Supplementary Tables

**Supplementary Table 1.** Number (and %) of *Klebsiella* genomes from the public (ENA) and GHRU genome collections that passed different quality control (QC) criteria.

| **QC criteria** | **Public genomes (total = 17,783)** | **GHRU genomes (total = 1,706)** |
|---|---|---|
| No. of contigs (≥1kb) <300 | 17,277 (97.2%) | 1,700 (99.6%) |
| N50 >25,000 | 17,538 (98.6%) | 1,706 (100%) |
| Genome length not >10% larger than largest genome and not <10% smaller than smallest genome in RefSeq of same species | 17,428 (98.0%) | 1,705 (99.9%) |
| Percentage contamination <5% (Confindr[1]) | 16,979 (95.5%) | 1,643 (96.3%) |
| Overrepresented sequences <1% (FastQC[2]) | 17,755 (99.8%) | 1,706 (100%) |
| Per-base sequence quality: a lower quartile that is <5 or a median of <20 for any base (FastQC) | 17,761 (99.9%) | 1,706 (100%) |
| Correct genus via Speciator (*Klebsiella* or *Raoultella*) | 17,686 (99.5%) | 1,706 (100%) |
| **All criteria met** | **16,537 (93.0%)** | **1,636 (96.0%)** |

**Supplementary Table 4**. Reference genomes used in the curation of the core gene set for *K. pneumoniae* analyses in Pathogenwatch.

| **Genome** | **ST** | **Reference** | **PMID** |
|---|---|---|---|
| NJST258_1 | 258 | DeLeo et al. 2014 | 24639510 |
| HS11286 | 11 | Liu et al. 2012 | 22408243 |
| MS6671 | 147 | Zowawi et al. 2015 | 26478520 |
| NTUH-K2044 | 23 | Wu et al. 2009 | 19447910 |
| PMK1 | 15 | Stoesser et al. 2014 | 25267672 |
| PittNDM01 | 14 | Doi et al. 2014 | 25070096 |
| EuSCAPE_PL046 | 3666 | David et al. 2020 | 32968015 |
| EuSCAPE_MT013 | 11 | David et al. 2020 | 32968015 |
| EuSCAPE_DE075 | 530 | David et al. 2020 | 32968015 |
| EuSCAPE_DE072 | 36 | David et al. 2020 | 32968015 |
| EuSCAPE_RS081 | 340 | David et al. 2020 | 32968015 |
| EuSCAPE_TR057 | 101 | David et al. 2020 | 32968015 |
| EuSCAPE_TR070 | 35 | David et al. 2020 | 32968015 |

**Supplementary Table 7**. Analysis tools available for different *Klebsiella* species in Pathogenwatch.

<table>
<tr><th>Species</th><th>MLST</th><th>cgMLST and core genome clustering</th><th>K- and O-loci identification</th><th>Resistance and virulence gene identification</th><th>Plasmid replicon identification</th><th>Tree-building</th></tr>
<tr><td>K. pneumoniae</td><td rowspan="5">Yes (K. pneumoniae scheme; https://bigsdb.pasteur.fr/klebsiella/)</td><td rowspan="5">Yes (K. pneumoniae scheme; https://bigsdb.pasteur.fr/klebsiella/)</td><td rowspan="17">Yes (Kleborate)</td><td rowspan="17">Yes (Kleborate)</td><td rowspan="17">Yes (Inctyper with Enterobacteriaceae database)</td><td>Yes</td></tr>
<tr><td>K. quasipneumoniae</td><td rowspan="16">NA</td></tr>
<tr><td>K. variicola</td></tr>
<tr><td>K. quasivariicola</td></tr>
<tr><td>K. africana</td></tr>
<tr><td>K. aerogenes</td><td>Yes (K. aerogenes scheme; https://pubmlst.org/organisms/klebsiella-aerogenes/)</td><td rowspan="12">NA</td></tr>
<tr><td>K. oxytoca</td><td rowspan="4">Yes (K. oxytoca scheme; https://pubmlst.org/organisms/klebsiella-oxytoca/)</td></tr>
<tr><td>K. michiganensis</td></tr>
<tr><td>K. grimontii</td></tr>
<tr><td>K. pasteurii</td></tr>
<tr><td>K. huaxiensis</td><td rowspan="7">NA</td></tr>
<tr><td>K. spallanzanii</td></tr>
<tr><td>K. (Raoultella) planticola</td></tr>
<tr><td>K. (Raoultella) ornithinolytica</td></tr>
<tr><td>K. (Raoultella) electrica</td></tr>
<tr><td>K. (Raoultella) terrigena</td></tr>
</table>

NA – not available.

**Supplementary Table 8.** *Klebsiella* species distribution among public (ENA) and GHRU genomes.

| **Species** | **Number (and %) of genomes in public collection** | **Number (and %) of genomes in GHRU collection** |
|---|---|---|
| *K. pneumoniae* | 14,635 (88.5%) | 1,451 (88.7%) |
| *K. quasipneumoniae* | 583 (3.5%) | 148 (9.0%) |
| *K. variicola* | 338 (2.0%) | 4 (0.2%) |
| *K. aerogenes* | 322 (1.9%) | 13 (0.8%) |
| *K. michiganensis* | 321 (1.9%) | 7 (0.4%) |
| *K. oxytoca* | 142 (0.9%) | 9 (0.6%) |
| *K. grimontii* | 99 (0.6%) | - |
| *K. (Raoultella) ornithinolytica* | 41 (0.2%) | - |
| *K. (Raoultella) planticola* | 20 (0.1%) | - |
| *K. quasivariicola* | 17 (0.1%) | 2 (0.1%) |
| *K. pasteurii* | 15 (0.1%) | 2 (0.1%) |
| *K. huaxiensis* | 3 (0.02%) | - |
| *K. (Raoultella) electrica* | 1 (0.01%) | - |
| **Total** | **16,537** | **1,636** |

**Supplementary Table 9.** Species assignments using automated laboratory methods and Speciator (Pathogenwatch) for 1,601 isolates identified as *Klebsiella* by both methods.

| | **Lab species ID** | | |
|---|---|---|---|
| **Speciator classification** | *K. oxytoca* | *K. pneumoniae* | *K. aerogenes* |
| *K. aerogenes* | | | 6 |
| *K. michiganensis* | 5 | 2 | |
| *K. oxytoca* | 7 | 2 | |
| *K. pasteurii* | 2 | | |
| *K. pneumoniae* | | 1,419 | 5 |
| *K. quasipneumoniae* | | 147 | |
| *K. quasivariicola* | | 2 | |
| *K. variicola* | | 4 | |
| **Total no. of isolates** | **14** | **1,576** | **11** |

**Supplementary Table 10**. Number (and %) of *K. pneumoniae* genomes from the GHRU and public (ENA) collections with each Kleborate virulence score (0 = none of the acquired virulence loci; 1 = yersiniabactin; 2 = yersiniabactin and colibactin, or colibactin only; 3 = aerobactin, without yersiniabactin or colibactin; 4 = aerobactin with yersiniabactin, without colibactin; 5 = yersiniabactin, colibactin and aerobactin).

| **Virulence Score** | **Colombia** | **India** | **Nigeria** | **Philippines** | **Total GHRU** | **ENA** | **Grand Total** |
|---|---|---|---|---|---|---|---|
| 0 | 237 (48.1) | 31 (10.1) | 94 (67.6) | 277 (54.1) | 639 (44.0) | 8127 (55.5) | 8766 (54.5) |
| 1 | 130 (26.4) | 159 (51.8) | 43 (30.9) | 213 (41.6) | 545 (37.6) | 4642 (31.7) | 5187 (32.2) |
| 2 | 122 (24.7) | 0 | 0 | 0 | 122 (8.4) | 960 (6.6) | 1082 (6.7) |
| 3 | 2 (0.4) | 0 | 2 (1.4) | 13 (2.5) | 17 (1.2) | 251 (1.7) | 268 (1.7) |
| 4 | 2 (0.4) | 117 (38.1) | 0 | 6 (1.2) | 125 (8.6) | 485 (3.3) | 610 (3.8) |
| 5 | 0 | 0 | 0 | 3 (0.6) | 3 (0.2) | 170 (1.2) | 173 (1.1) |
| Total | 493 (100) | 307 (100) | 139 (100) | 512 (100) | 1451 (100) | 14635 (100) | 16086 (100) |

### Supplementary Figures

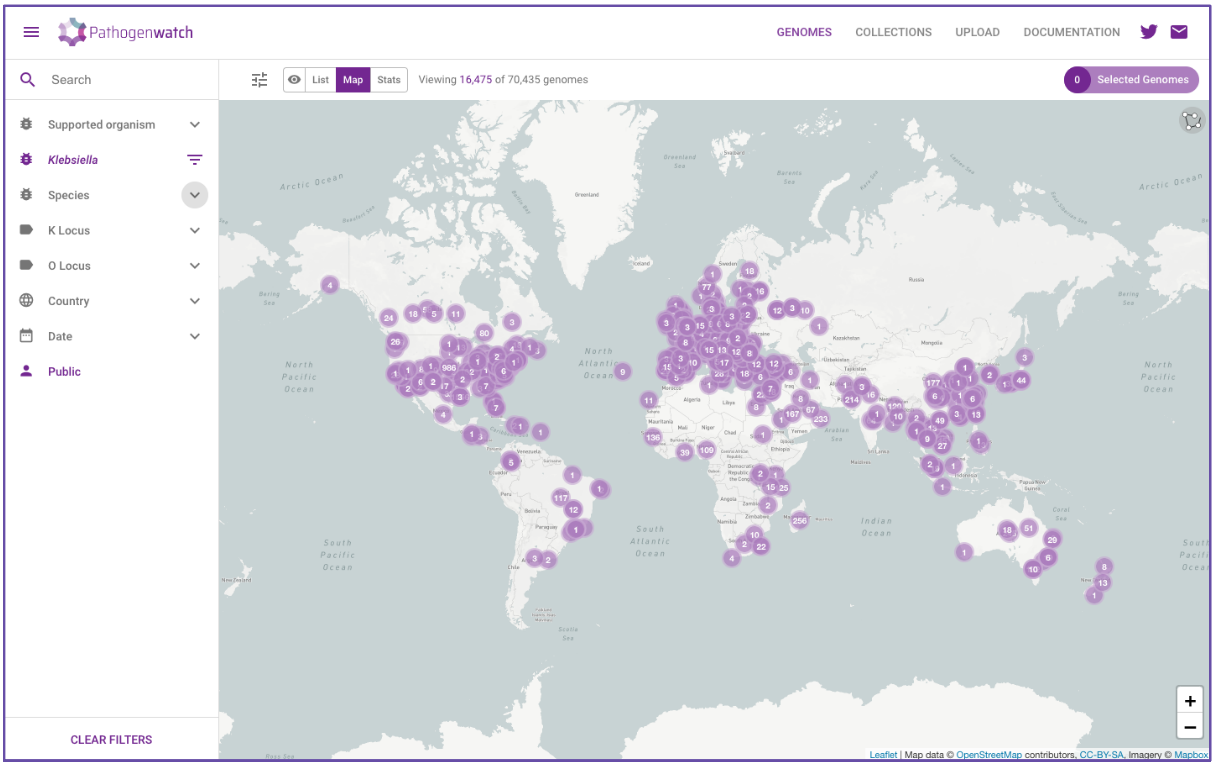


**Supplementary Figure 1.** Screenshot of the Pathogenwatch application showing the distribution of sampling locations of all public *Klebsiella* genomes.

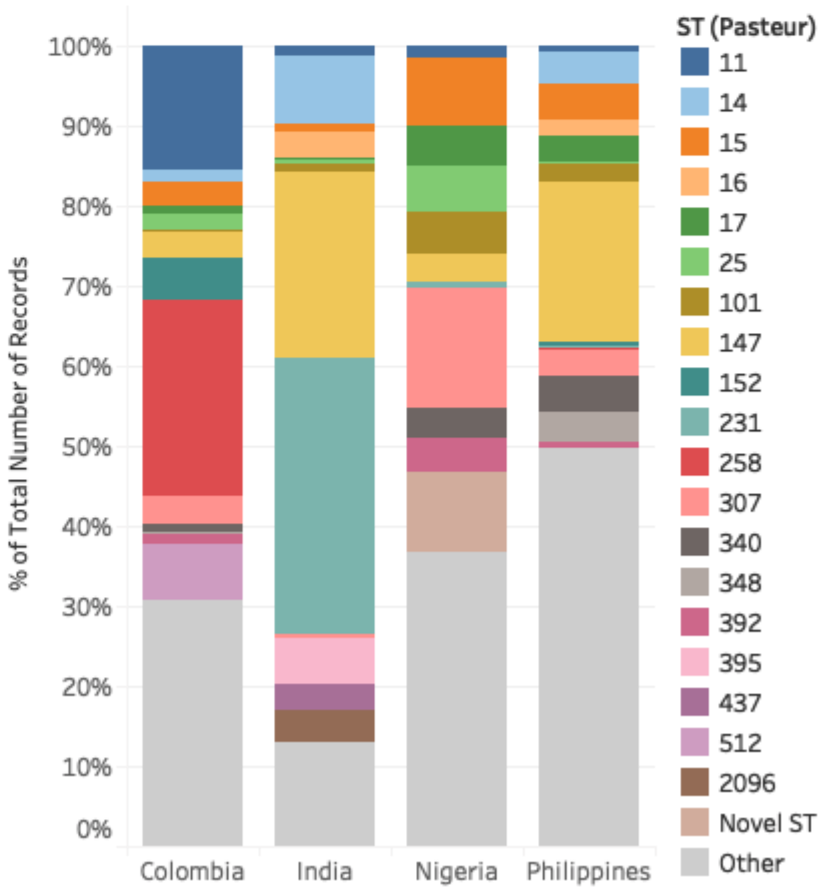


**Supplementary Figure 2.** Relative frequency of sequence types (STs) among *K. pneumoniae* genomes from the GHRU participant countries. The combined top seven STs from each country are shown, while the remaining 189 STs are grouped as "Other".

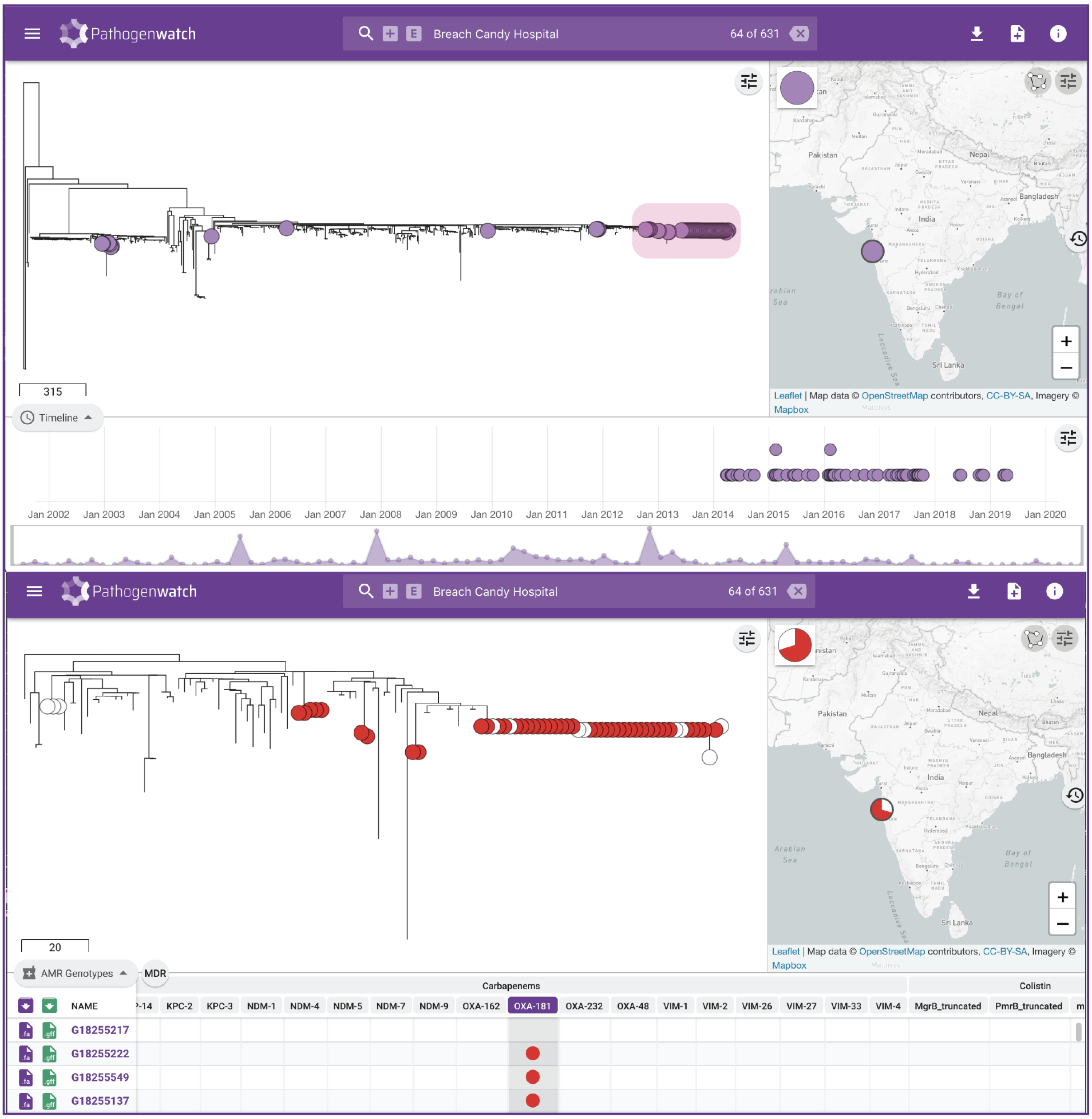


**Supplementary Figure 3.** Application of Pathogenwatch to local outbreak investigations. The top panel shows a phylogenetic tree of 631 ST147 *K. pneumoniae* genomes from the global and GHRU collections. The purple tree nodes indicate genomes from a hospital in India (see map) and their time of isolation (see timeline). The clade highlighted in the pink shaded box is shown in detail in the bottom panel. Only nodes from the hospital are shown, with red nodes additionally indicating the presence of carbapenemase gene OXA-181.

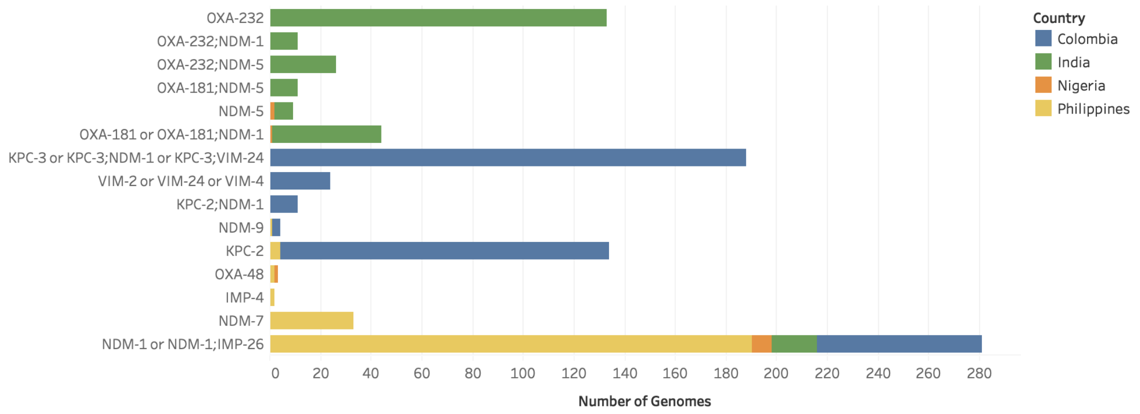


**Supplementary Figure 4.** Distribution of carbapenemase genes among 914 *K. pneumoniae* genomes from the GHRU participant countries.

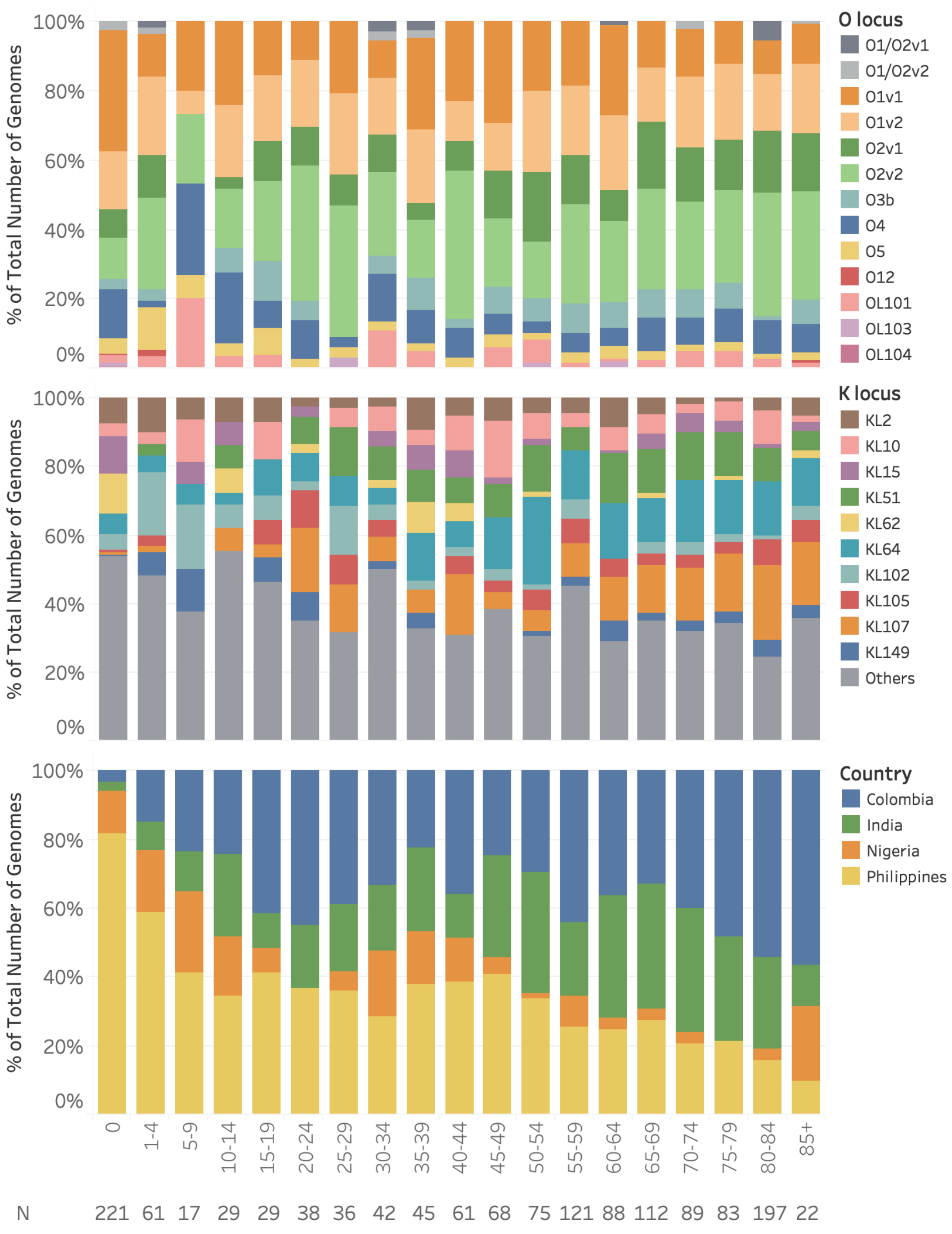


**Supplementary Figure 5.** Distribution of O- and K-locus types in *K. pneumoniae* isolates from the GHRU collection stratified by age. All the predicted O-types (top panel) and the ten most frequent K-types (middle panel) are shown. Other K-types are shown in grey. The distribution of isolates from each GHRU participant country is shown in the bottom panel. The number of isolates in each age category are shown below the x-axis (N).

2. FastQC. https://www.bioinformatics.babraham.ac.uk/projects/fastqc/
